# Chaperonin Abundance Boosts Bacterial Fitness

**DOI:** 10.1101/2019.12.31.891820

**Authors:** C. M. Santosh Kumar, Kritika Chugh, Anirban Dutta, Vishnuvardhan Mahamkali, Tungadri Bose, Sharmila S. Mande, Shekhar C. Mande, Peter A. Lund

**Affiliations:** School of Biosciences and Institute of Microbiology and Infection, University of Birmingham, B15 2TT, UK; Department of Biotechnology and Bioinformatics, University of Rajasthan, Jaipur – 302004, India; Bio-Sciences R&D Division, TCS Research, Tata Consultancy Services Limited, Pune - 411013 India; Australian Institute for Bioengineering and Nanotechnology (AIBN), The University of Queensland, Brisbane - 4072, Australia; Laboratory of Structural Biology, National Centre for Cell Science (NCCS), Pune - 411027, India

**Author notes:** Address Correspondence to C. M. Santosh Kumar.

**Keywords:** Metabolic flux, Chaperonin, GroEL, Evolution, Proteomics, Metabolism

## Abstract

The ability of chaperonins to buffer mutations that affect protein folding pathways suggests that their abundance should be evolutionarily advantageous. Here, we investigate the effect of chaperonin overproduction on cellular fitness in *Escherichia coli*. We demonstrate that chaperonin abundance confers (a) an ability to tolerate higher temperatures, (b) improved cellular fitness and (c) enhanced folding of metabolic enzymes, which is expected to lead to enhanced energy harvesting potential.

## Introduction

Chaperonins are found in nearly every organism across all domains of life, and are essential in all cases tested to date, although in some cases non-essential paralogues are found (Kumar, 2017; Lund, 2009). The GroE chaperonin system of *E. coli*, consisting of the 60 kDa GroEL and the 10 kDa GroES proteins assembled into ring complexes of 14 and 7 sub-units, respectively, is encoded by the *groE* operon (Balchin et al., 2016; Bukau and Horwich, 1998). This operon is expressed principally from two promoters, one utilized in the presence of housekeeping sigma factor σ^70^, and the other, which is strongly induced due to the accumulation of unfolded proteins, in the presence of the alternative sigma factor, σ^32^ (RpoH) (Kumar et al., 2015; Lund, 2001; Schumann, 2017). As σ^32^ levels respond to unfolded protein, this provides a feedback loop to maintain proteostasis (Kim et al., 2013). When cells are shifted to heat shock temperatures between 42 – 46 °C, GroE levels increase by 5 – 10 fold, reaching up to 12% of the entire cellular proteome (Martin et al., 1992). These increased levels interact more extensively with the proteome and are assumed to prevent misfolding or assist refolding of heat-stressed proteins (Houry et al., 1999; Llorca et al., 1998; Martin et al., 1992). Cells that cannot mount an unfolded protein response due to *rpoH* deletion are extremely temperature sensitive, and selection for pseudo-revertants of these strains at elevated temperatures yields up-promoter mutations in the *groE* promoter (Kusukawa and Yura, 1988). GroE is thus important even under normal growth conditions, and indeed GroEL and GroES are respectively the 20^th^ and 21^st^ most abundant proteins in *E. coli* (excluding ribosomal proteins), with sufficient protein being made under non-stressed conditions to produce approximately 2800 complexes of GroEL and 5700 complexes of GroES (Li et al., 2014). Other chaperones that are also abundant include the ribosome bound trigger factor (TF), which is the 19^th^, and the Hsp70 homologue, DnaK, which is the 27^th^ most abundant. The high levels of all these chaperones indicates their key roles in cell growth. Although combined loss of TF and DnaK is deleterious to cells, GroEL and GroES are the only chaperones in *E. coli* that are essential under all conditions (Fayet et al., 1989).

GroE assists the folding of 10-15% cellular proteins (Houry et al., 1999), many of which are essential (Kerner et al., 2005). GroE’s ability to fold ‘folding-compromised’ proteins (Fares et al., 2002; Houry et al., 1999; Kerner et al., 2005; Tokuriki and Tawfik, 2009) is consistent with a ‘genetic capacitance’ function. Many studies with different heterologous proteins have shown that GroE can enhance their folding (Durao et al., 2015; Ishimoto et al., 2014; Tokuriki et al., 2008; Tokuriki and Tawfik, 2009; Wyganowski et al., 2013). In addition, some deleterious mutations are retained in the genome upon overexpression of *groE*, probably due to chaperonin-buffered folding of polypeptides whose folding pathway has been perturbed (Fares et al., 2002; Sabater-Munoz et al., 2015; Van Dyk et al., 1989; Williams and Fares, 2010). However, since GroE is an active ATPase, its overproduction could be deleterious. In this study, we have assessed the effect of GroE overproduction on the growth characteristics and thermal tolerance of *E. coli*, and used proteomics and *in silico* flux balance analysis to determine the likely impact of chaperonin overproduction on the metabolic advantage and consequent fitness of the organism.

## Results

### Construction of GroE Overproducing Strains

To investigate the effect of chaperonin overproduction on *E. coli*, we constructed two chaperonin producing strains, GL-H_t_ and GL-L_t_, which produce high and low levels of GroE (Fig. 1A). These strains were derived from strain *E. coli* LG6 (Horwich et al., 1993), in which the *P*_*groE*_ promoter is replaced by the *P*_*lac*_ promoter, by transforming with pTrc-GSL, which overexpress *groE* operon upon induction with lactose, or its parental plasmid pTrc99A. SDS-PAGE confirmed significant overproduction of GroEL (Fig. 1B) and GroES (Fig. 1C) in GL-H_t_ compared to GL-L_t_. From Western blotting of the lysates we estimate that GroEL levels are twenty-fold greater in GL-H_t_ than in GL-L_t_ (Fig. 1D). The expression levels of GroEL in GL-L_t_ were lower than the MG1655, where wildtype *P*_*groE*_ promoter drives the expression (Fig. 1B-D) (Chapman et al., 2006).

**Figure 1.**
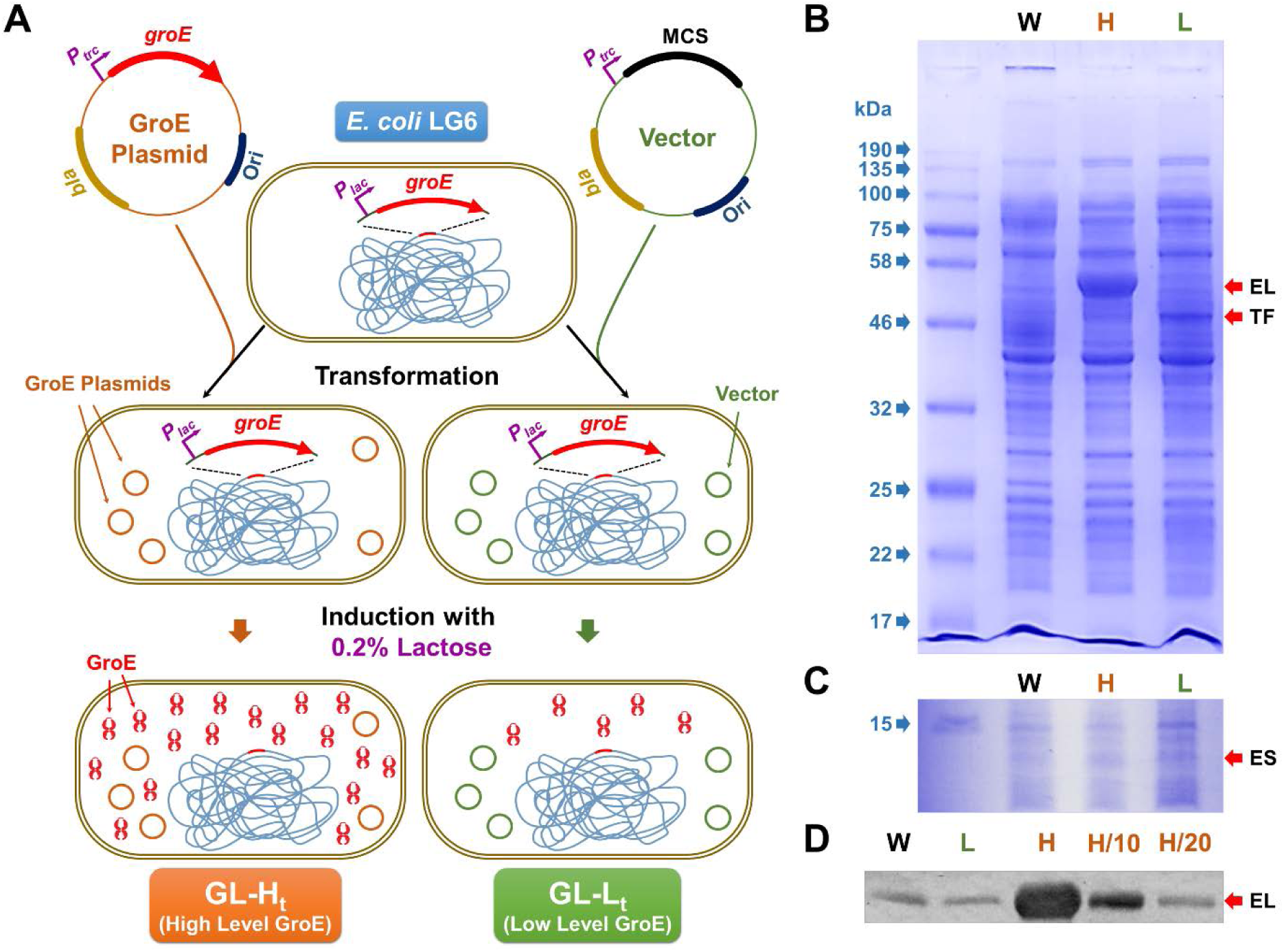
A. Construction of GL-L_t_ and GL-H_t_ Strains. In *E. coli* LG6, the *groE* operon is under the control of the inducible *P*_*trc*_ promoter. This strain was transformed with either a plasmid expressing *groE* under the control of lactose (GroE Plasmid), pTrc-GSL or its empty vector (Vector), pTrc99a. Upon culturing in the presence of lactose, GL-H_t_ produces elevated levels of GroEL and GroES, due to the induction of chromosomal and plasmid-borne *groE* operon, while GL-L_t_ will have lower production of GroES and GroEL due to the induction of only the chromosomal *groE* copy. **B and C. SDS-PAGE Confirms Enhanced Levels of GroE in GL-H**_**t**_. 12.5% (**B**) and 20% (**C**) SDS-PAGE gels comparing the levels of GroEL (EL) and GroES (ES), respectively, in GL-H_t_ (H), GL-L_t_ (L) and the wildtype *E. coli* MG1655 (W) strains. Numbers and arrows in blue indicate molecular masses of standards. TF indicates trigger factor. **C. Immunoblotting Estimates Enhancement of GroEL in GL-Ht**. Lysates of MG1655 (W), GL-L_t_ (L) and GL-L_t_ (H) were resolved on a 12.5% SDS-PAGE and probed with GroEL specific antibody. H/10 and H/20 indicate the lanes with ten and twenty fold diluted GL-H_t_ lysates, respectively.

### GroEL-GroES Overproducing Strains showed Enhanced Temperature Tolerance

As GroE is involved in protection against thermal stress, we analyzed the impact of different GroE levels in GL-H_t_ and GL-L_t_ on growth at temperatures ranging from 17°C to 48°C (Fig. 2) (Chilukoti et al., 2015). *E. coli* MG1655 and MG1655 hosting pTrc-GSL were included for comparison. As expected, GL-L_t_ cells exhibited heat and cold sensitive phenotypes and consequently showed poor growth at many temperatures, consistent with previous observations that sufficient levels of GroES-GroEL are required for growth over a wide temperature range (Ferrer et al., 2003). Further, MG1655 showed much better temperature tolerance than GL-L_t_, showing the importance of the heat-shock regulation of the *P*_*groE*_ promoter. The strains harboring pTrc-GSL tolerated higher temperatures, up to 48 °C, than the vector-only MG1655, where *groE* expression is temperature regulated, suggesting that higher levels of GroEL-GroES enable higher temperature tolerance.

**Figure 2.**
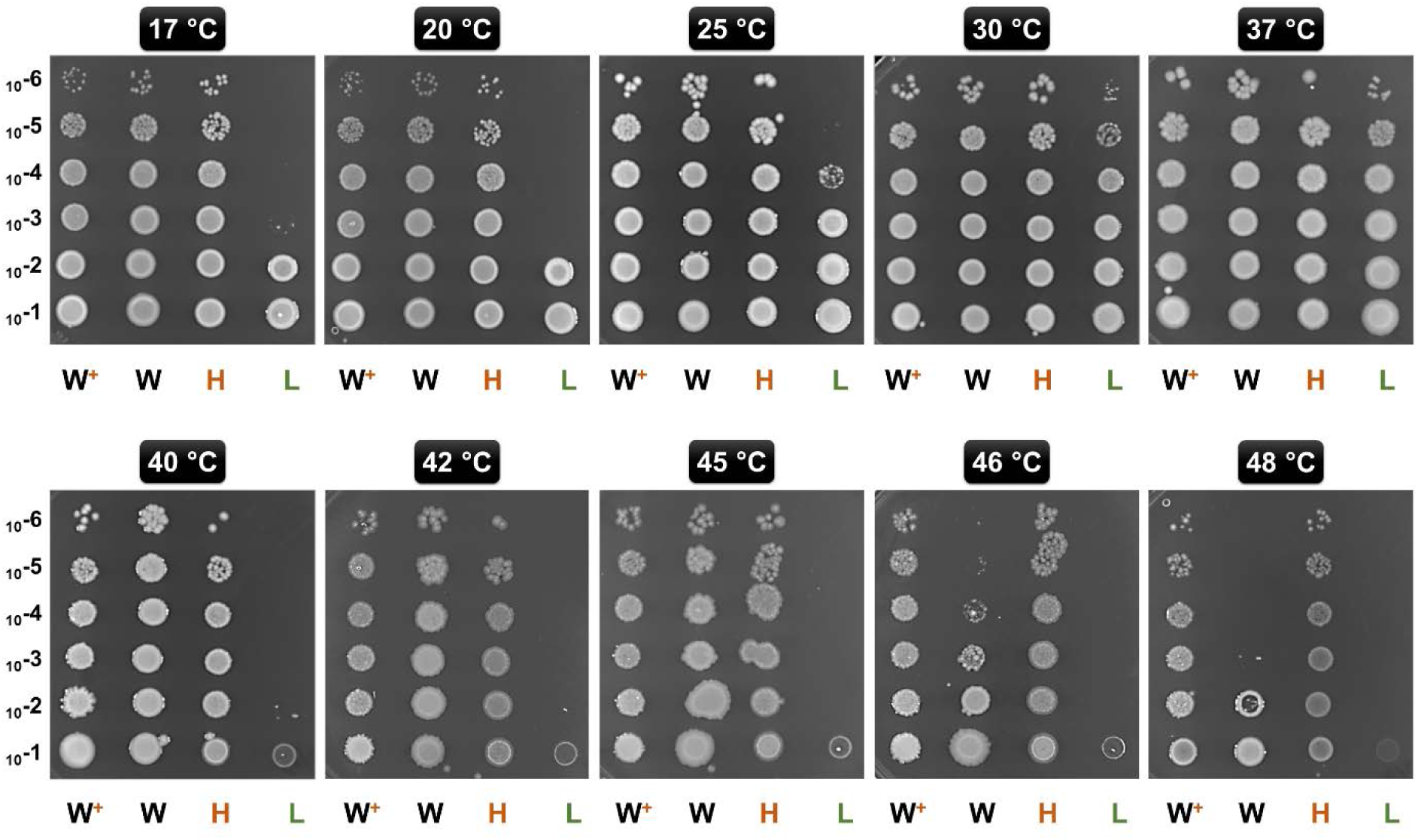
Temperature Tolerance upon Chaperonin Overproduction. Ten-fold serially diluted (10^−1^ through 10^−6^) exponentially growing cultures of GL-H_t_ (H), GL-L_t_ (L), MG16155 (W) and MG1655 with pTrc-GSL (W^+^) were spotted onto LB agar plates supplemented with lactose. These plates were incubated at the indicated temperatures.

### GroEL-GroES Overproducing Strain Exhibited Fitness Advantage in Competition Culture

Since higher levels of chaperonins led to a growth advantage, we examined whether this translated to a fitness advantage even under low stress conditions, by competing two strains with different GroE levels. Since two strains showed similar growth profiles on the plates and in independent liquid cultures at 30 °C (Fig. S2), we chose this temperature for the competition culture. To do these experiments, we needed to be able to control the plasmid borne and chromosomal copies of the *groESL* operon independently. Therefore, we constructed two new strains with a *P*_*BAD*_ based plasmid expression system, called GL-H_b_ (high expression) and GL-L_b_ (low expression) strains. Similar to GL-H_t_, GL-H_b_ showed several fold higher GroE induction levels (Fig. S1A) and temperature resistance (Fig. S1B). The cultures of GL-H_b_ and GL-L_b_ were competed for 20 passages (∼700 generations) and their relative fitness(s) were estimated (Fig. 3A) as described in Materials and Methods (Macho et al., 2010; Monk et al., 2008; van Opijnen and Camilli, 2013). The high *groE* expressing GL-H_b_ outcompeted GL-L_b_ (Fig. 3B), indicating that chaperonin level is an important fitness determinant.

**Figure 3.**
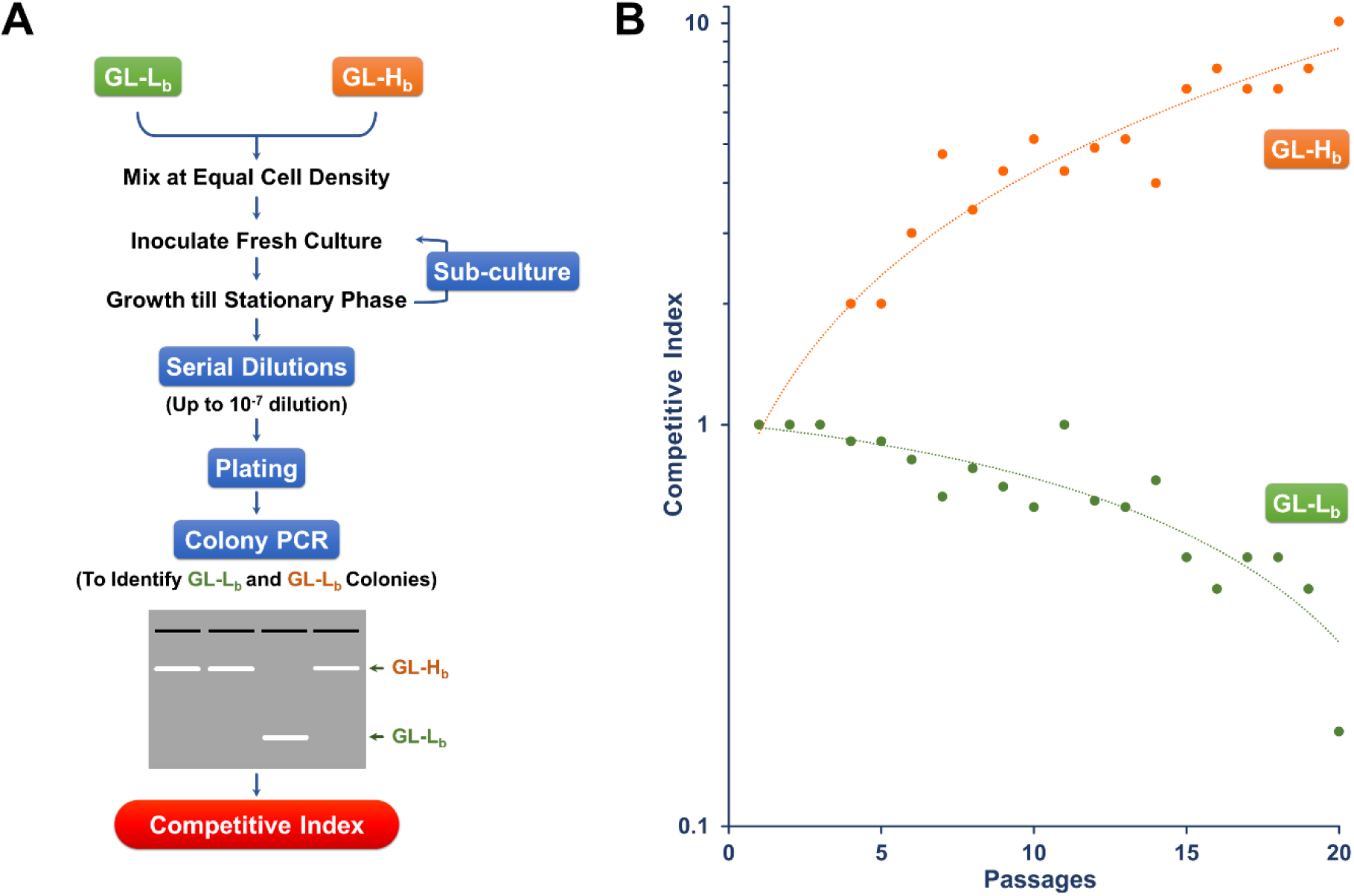
Chaperonin Depletion leads to Lower CI. **A.** Strategy for Determining the CI for GL-L_b_ and GL-H_b_ strains. Stationary phase cultures of GL-L_b_ and GL-H_b_ strains were mixed at equal cell density, grown to stationary phase and sub-cultured in fresh media for 20 continuous passages. Cells recovered at each passage were serially diluted as indicated, spread on LB agar plates and the resulting colonies were scored for their phenotype (GL-L_b_ or GL-H_b_), by colony PCR, using vector specific oligonucleotide primers. **B.** CI, as a degree of fitness, was calculated at every passage from a ratio of proportion of the cells with a particular phenotype and plotted as a function of number of passages. CI trend was similar among the three independent experiments.

### Proteomic Analysis Reveals Preferential Enrichment of Metabolic Enzymes in GroEL-GroES Overproducing Strains

Overproduction of a chaperonin is likely to enrich the levels of folded proteins in the cells, while unfolded or misfolded proteins tend to remain insoluble and thereby targeted to either the inclusion bodies or marked for degradation (Samuelson, 2011). Given this context, we investigated the proteomes of GL-H_b_ and GL-L_b_ cells, to identify what might account for the differences in fitness. Both strains were grown under identical conditions (Fig S2) and their soluble proteome profiles (on 2D PAGE) were compared for relative abundance (Fig. S3 and Table S1). Although many of the identified proteins were known chaperonin clients of classes I – III (Kerner et al., 2005), class IV (Arora et al., 2016) and class V (Chapman et al., 2006), several proteins identified as being differentially expressed were not known clients (Table S1), suggesting that either chaperonin overexpression can indirectly affect the folding of these non-client proteins or that the chaperonin client base is larger than currently understood.

Interestingly, none of the obligate GroEL clients – class III (Kerner et al., 2005) were enriched in GL-L_b_ strain, suggesting that the lowered chaperonin level is insufficient to fold these proteins. Notably, the outer membrane proteins, OmpC and OmpF, which are involved in metabolite import and are known GroE clients (Kerner et al., 2005), were enriched in the soluble proteome of GL-L_b_. Consequently, these proteins were less abundant in the membrane fractions of GroEL-deficient strains (Fig. S5). Further, the enrichment of TF in GroEL-deficient GL-L_b_ (Fig. 1B and Table S1), suggested compensatory functions for low GroE. Furthermore, enrichment of several metabolic enzymes in GL-H_b_ strain, suggested a higher rate of metabolism in this strain. To evaluate this hypothesis, we collated publicly available *E. coli* proteomic data from the paxdb database (Wang et al., 2012), screened for proteins that were co-enriched with GroE across different experiments and identified 404 proteins that showed significant correlation, in expression levels, with GroE (Pearson correlation co-efficient > = 0.7, *P* < 0.05). Interestingly, a GO enrichment analysis of this set of proteins revealed that majority of these proteins were involved in metabolism and energy production, including multiple GO terms related to carbohydrate metabolism (Table S3).

### Flux Balance Analysis of oxidative phosphorylation in high- and low-GroEL strains

The metabolic FBA simulations analysis was carried out using ‘TransFlux’, an in-house tool with a module to incorporate gene expression/proteomic profiles in the FBA framework. The proteomic profiles (summarized in supplemental materials and Table S1) and observations from *E. coli* gene expression microarray studies, derived from the Many Microbe Microarrays Database (M3D, www.m3d.mssm.edu) (Faith et al., 2008) were utilized to constrain fluxes though respective reactions, while performing two independent FBA simulations, each of which corresponded to the expression/enrichment profiles of the enzymes enriched in GL-L_b_ and GL-H_b_ strains. As expected, higher flux was observed through several pathways of carbon metabolism including glycolysis, gluconeogenesis, citric acid cycle (TCA cycle) and its anaplerotic reactions, and alternate carbon metabolism, in the simulated GL-H_b_ strain (Table S2). Apparently, these pathways are supported by enhanced import of glucose and glycerol (Supplementary File 2). Further, pathways corresponding to several glucogenic amino acids metabolism and energy generating oxidative phosphorylation were enriched in this strain. However, the pathways leading to the toxic Methylglyoxal synthesis have been enriched in the GL-H_b_ strain (Table S2). However, pathways leading to the metabolism of membrane lipids, pyruvic acid, pentose sugars, ubiquinone and salvage of nucleotides are enriched in the GL-L_b_ strain. Overall, FBA simulations indicated that the metabolic enzymes that were enriched in GL-H_b_ potentially lead to higher metabolic flux in this strain (Table S2 and Supplementary material). Further inadequate respiration was observed in GL-L_b_ (Table S2). Consistent with this analysis, the GroEL-deficient strain showed reduced growth relative to the high GroEL strain under low oxygen conditions (Fig. S4).

## Discussion

Over- or under-production of chaperonins in several organisms has been demonstrated to perturb rates of proteolysis (Martinez-Alonso et al., 2010), influence growth rates, and alter the expression levels of compensatory chaperones like DnaK (Lemos et al., 2007). Here we present a simple model system to study the effects of GroE overproduction (Fig. 1). We demonstrate that the overexpression of GroE chaperonin results in enhanced thermal tolerance (Fig. 2) and competitive advantage (Fig. 3). GroEL is known to be required for growth at low (Ferrer et al., 2003) and high (Guisbert et al., 2004) temperatures. Consistent with this, the GroEL-deficient strains exhibited both cold and heat sensitive phenotypes (Fig. 2) Proteomic studies (Table S1) followed by flux balance analysis (Supplementary material) suggest that the acquired fitness advantage could be attributed to an enriched set of metabolic enzymes. Chaperonin depletion was observed to induce the enrichment of the compensatory chaperone, TF (Fig. 1B and Table S1), which may act as a *holdase* for the GroE client proteins (Hartl and Hayer-Hartl, 2002). Interestingly, while GroE is more abundant than TF in *E. coli* (Zou et al., 2014), TF is observed to be abundant in mycoplasma which lack *groE* operon (Bang et al., 2000; Lund, 2009; Musatovova et al., 2006; Weiner et al., 2003), suggesting that higher levels of TF might be needed in such bacteria to compensate for the chaperonin deficiency. The TF – GroEL interplay, owing to their overlapping functions and client-base (Avellaneda et al., 2017; Bhandari and Houry, 2015), has been demonstrated both *in vitro* (Kandror et al., 1995) and *in vivo* in *E. coli* (Kandror et al., 1997). Therefore, it seems likely that TF enrichment in GL-L_b_ is compensating for GroE depletion and that TF may be holding the substrates (Singhal et al., 2015). Further, the enrichment of the outer-membrane proteins OmpC and OmpF in the soluble proteome of GL-L_b_ suggests that these known GroEL client proteins failed to reach their final cellular destinations and may have remained soluble, possibly in a TF-bound state. The reduced levels of these metabolite transporters in the membranes of GroEL deficient strain (Fig. S5) might be responsible, in part, for the lower metabolite transport and metabolic flux in this strain (Table S2). The upregulation of TF was not observed in MG1655, even though GroE expression levels were lower, which may be because in the wild-type strain, levels of GroE respond directly to levels of unfolded proteins. The higher fitness of the GroES and GroEL over-producing strains under the conditions of our experiments is likely to be associated with fitness costs under other conditions (Fig. 2 and 3), otherwise it would be expected that higher expression would have evolved. In this regard, it is interesting to note that growth in the absence of aeration was reduced in GroEL-GroES over-producing strains (Fig. S2).

This study shows a direct relation between abundance of chaperonins and competitive fitness, which begs the question as to why evolution has not selected for strains with higher levels of chaperonin producing capability. The predictions from FBA simulations provide some clues that may answer this. Although enhanced glycolysis, TCA cycle and oxidative phosphorylation in the GL-H_b_ cells increase cellular energy currency, FBA simulations for the GL-H_b_ strain also predicted an enhanced production of a toxic side product, methylglyoxal (Table S2), a very toxic three-carbon aldehyde that can inhibit *E. coli* growth at millimolar concentrations (Kayser et al., 2005; Weber et al., 2005). Therefore, evolution might have selected a balance in metabolic states between energy production and methylglyoxal toxicity, which would have, in turn, selected for an optimal level of chaperonin expression. The fact that chaperonins are active ATPases provides another possible answer to this question. Overabundance of chaperonins might be linked to ATP depletion and consequent reduced growth (Sabater-Munoz et al., 2015). Thus high levels of chaperonin expression would have been selected against during the course of evolution. These explanations are not exhaustive and the final level of chaperonin expression selected for is likely to result from a balance of optimizing fitness due to a multitude of factors.

Our analysis showed that GroE over-production results in several pleiotropic consequences that can enhance cellular fitness under certain conditions. These observations need to be probed further, using tools to enhance our understanding of the precise role of the chaperone-client interaction in influencing fitness and, ultimately, evolution. A similar system could be advantageous in studying the effect of chaperonin overproduction in different microbes, especially those with multiple chaperonins (Kumar, 2017).

## Materials and Methods

### Materials, Plasmids, Bacterial Strains and Growth Conditions

All chemicals were from Sigma, Inc. Bacterial growth media and media supplements were from HiMedia Laboratories, Inc., Mumbai, India. Phusion polymerase for colony PCR was purchased from New England Biolabs Inc, USA. GroE expression plasmids, pBAD-GSL and pTrc-GSL were generated by cloning GroE operon into NcoI and HindIII sites on plasmids pBAD24 (Guzman et al., 1995) and pTrc99A (Amann et al., 1988), respectively. The *groE* conditional mutant strain, *E. coli* LG6, was a kind gift from Arthur Horwich, Yale University, USA (Horwich et al., 1993). This strain produces GroE at levels similar to the wildtype at 30 °C (Chapman et al., 2006). Oligonucleotide primers were purchased from Integrated DNA Technologies, Inc., Coralville, IA, USA.

### Construction and validation of strains producing high and low GroE levels

To enable control of GroE levels independently from the growth temperature, two strains that differentially express *groE* were generated from the *E. coli* strain LG6, in which the chromosomal *groE* promoter has been replaced with a *P*_*lac*_ promoter (Horwich et al., 1993). A high level GroE expression strain, GL-H_t_ (for <u>G</u>roE<u>L H</u>igh p<u>T</u>rc), was obtained by transforming LG6 with pTrc-GSL and a lower level GroE expression strain, GL-L_t_ (for <u>G</u>roE<u>L L</u>ow p<u>T</u>rc) was obtained by transforming with the control plasmid pTrc99A (Amann et al., 1988). The scheme for the generation of these phenotypes is illustrated in Fig. 1A. To confirm the expression levels, these strains were cultured in the presence of 0.2% D-lactose to induce chromosome and plasmid borne *groE* operons, for 3 h at 30 °C. The resulting cells were suspended in lysis buffer containing 50 mM HEPES:KOH pH 7.5 and 150 mM NaCl, 1 mM EDTA and 1 mM PMSF, mixed with Lysing Matrix E and lysed by homogenization in FastPrep (M. P. Biomedicals, Irvine, CA, USA). Lysates were centrifuged at 13,000 rpm for 20 min to obtain soluble lysates. The soluble lysates were resolved on 12.5% and 20% SDS-PAGE followed by Coomassie Brilliant Blue staining to detect the levels of GroEL and GroES, respectively. In parallel, these lysates were probed with an anti-GroEL monoclonal antibody (1.10B) at 1:100 dilution and the blots were developed by BCIP/NBT-Purple Liquid Substrate System (Sigma Aldrich Inc., St. Louis, MO, USA). In addition to these strains, two strains that enable independent regulation of the chromosome and plasmid borne copies of *groE* operon were generated by transforming LG6 with pBAD-GSL and pBAD24 to result in high and low expression strains, GL-H_b_ and GL-L_b_, respectively. These strains were cultured in the presence of 0.2% lactose plus 0.2% arabinose to obtain the high and low expression levels (Fig. S1A).

### Temperature Sensitivity Assessment

To assess the extent to which GroE overproduction enables temperature tolerance, actively growing cultures of GL-H_t_ and GL-L_t_ were normalized for OD_600_, serially diluted, and spotted onto eight LB agar plates supplemented with 0.2% D-lactose. The plates were incubated at 17 °C, 20 °C, 22 °C, 25 °C, 30 °C, 37 °C, 40 °C, 42 °C, 45 °C, 46 °C and 48°C. Wildtype MG1655 harboring pTrc-GSL or empty vector (pTrc99A), respectively, were included as controls.

### Competition and estimation of relative fitness

GL-H_b_ and GL-L_b_ cells were subjected to competitive serial culturing as described previously (Smith, 2011; Vulic and Kolter, 2001; Zambrano et al., 1993). Briefly, equal number of cells from these two cultures were mixed and grown in fresh LB supplemented with 0.2% L-arabinose and 0.2% D-lactose. This mixed culture was grown to stationary phase at 30 °C, recovered, labelled Passage-1 and used to generate the second passage (Fig. 2A). Serial sub-culturing was repeated for further 20 passages (∼700 generations). At each passage, a fraction of the cultures was serially diluted up to 10^−7^ dilution in LB broth and spread on LB agar plates supplemented with 0.2% D-lactose, which supports the growth of the cells derived from either strain. The resulting colonies at each passage, in the range of 23-28 colonies, were screened using colony PCR to identify whether colonies were derived from either GL-H_b_ or GL-L_b_ cells. Colony PCR with the PBADF (5’-CTGTTTCTCCATACCCGTT-3’) and PBADR (5’-CTCATCCGCCAAAACAG-3’) primers, which bind upstream and downstream of the MCS on the parental vector pBAD24, results in the amplification of 2.1 and 0.3 kb fragments from the pBAD-GSL and pBAD24 vectors, harbored by the GL-H_b_ and GL-L_b_ cells, respectively. Relative competitive index (CI), a measure of relative fitness, was calculated for each phenotype as the ratio of the proportion of a particular cell type at the final and initial generations (Macho et al., 2010; Monk et al., 2008; van Opijnen and Camilli, 2013).

### Proteomic Analysis

Equal number of cells from exponentially growing cultures (OD_600_ = ∼ 0.6) of GL-H_b_ or GL-L_b_ strains were harvested, suspended in lysis buffer (50 mM HEPES:KOH pH: 7.5 and 150 mM NaCl, 1 mM EDTA and 1 mM PMSF), lysed by sonication, and the soluble protein fractions were recovered by centrifugation at 12,000 rpm for 20 min. 200 µg protein from the soluble fractions of each lysate were resolved through 2D PAGE following the standard protocols. Briefly, the lysates were resolved on the first dimension through a 7 cm Immobilized pH Gradient (IPG) strip of 3 – 10 pH range, followed by 10% SDS-PAGE on the second dimension. The separated proteins were stained with Coomassie brilliant blue and intensities of the stained protein spots were compared between the two gels using densitometry. Differentially enriched spots between the two lysates were picked and identified by tandem mass-spectrometry in an LTQ Orbitrap Mass Spectrometer (Thermo Fisher Scientific Inc., Waltham, MA, USA). The differentially enriched proteins were identified using MASCOT (Hirosawa et al., 1993) search against UniProtKB/TrEMBL (UniProt, 2019) and RefSeq (O’Leary et al., 2016) databases. The spot identification was done in collaboration with the Centre for Cellular and Molecular Platforms, Bangalore, India.

### Flux Balance Analysis of the GL-H_b_ and GL-L_b_ Strains

*E. coli* genome-scale metabolic network iJO1366 (Blais et al., 2013) was used for performing the FBA simulations. The iJO1366 model was first simulated using a standard energy source (equivalent of a glucose-supplemented minimal media) to obtain the steady state fluxes through each of the reactions (Orth et al., 2011). The objective function of this FBA simulation was to maximize the biomass production, while using some ‘default constraints’ (lower- and upper-bounds of fluxes through each reaction) derived from the literature (Blais et al., 2013). Following this preliminary assessment of the *E. coli* cell’s metabolic potential, two independent FBA simulations were performed, each of which corresponded to the enzyme expression/enrichment profiles of the GL-H_b_ and GL-L_b_ strains. During each of these simulations the reaction flux values were appropriately constrained, based on the results from the preliminary assessment and the corresponding enzyme expression/enrichment profiles (Supplementary File 2). Incorporating enzyme expression profiles into FBA simulations was performed with our software tool ‘TransFlux’, developed in-house and housed at http://www.nccs.res.in/TrasFlux/index.jsp. Details of the parameters and the principles applied in FBA are presented in the supplementary methods section.

## Acknowledgements

We would like to acknowledge Biotechnology and Biological Sciences Research Council, the Royal Society, British Academy and Academy of Medical Sciences, UK for funding. We would like to thank, Arthur Horwich for the GroE depletion strain, *E. coli* LG6 and cCAMP, Bangalore for assistance in proteomic studies. We thank Abhijit Sardesai and Gaurang Mahajan for helpful discussions, Melanie Swannell, Amanda Rossiter, Anna Schager, Chistopher Icke, Nitin Bayal, Sapna Sugandhi and Surbhi Dhingra for support in the initial studies, and Ishita Verma for helping in designing the TransFlux website. Sharmila S. Mande, Anirban Dutta and Tungadri Bose are employees of TCS Research (Tata Consultancy Services Ltd., Pune, India), and would like to acknowledge TCS for its support. Initial part of this work was supported by grants from the Department of Biotechnology, India (BT/PR3260/BRB/10/967/2011). The authors declare that no competing financial interests exist.

## References

Amann, E., Ochs, B., and Abel, K.J. (1988). Tightly regulated tac promoter vectors useful for the expression of unfused and fused proteins in Escherichia coli. Gene 69, 301–315.

Arora, G., Sajid, A., Virmani, R., Singhal, A., Kumar, C.M.S., Dhasmana, N., Khanna, T., Maji, A., Misra, R., Molle, V., et al. (2016). Ser/Thr protein kinase PrkC-mediated regulation of GroEL is critical for biofilm formation in Bacillus anthracis. npj Biofilms and Microbiomes 3, 7.

Avellaneda, M.J., Koers, E.J., Naqvi, M.M., and Tans, S.J. (2017). The chaperone toolbox at the single-molecule level: From clamping to confining. Protein Sci 26, 1291–1302.

Balchin, D., Hayer-Hartl, M., and Hartl, F.U. (2016). In vivo aspects of protein folding and quality control. Science 353, aac4354.

Bang, H., Pecht, A., Raddatz, G., Scior, T., Solbach, W., Brune, K., and Pahl, A. (2000). Prolyl isomerases in a minimal cell. Catalysis of protein folding by trigger factor from Mycoplasma genitalium. Eur J Biochem 267, 3270–3280.

Bhandari, V., and Houry, W.A. (2015). Substrate Interaction Networks of the Escherichia coli Chaperones: Trigger Factor, DnaK and GroEL. Adv Exp Med Biol 883, 271–294.

Blais, E.M., Chavali, A.K., and Papin, J.A. (2013). Linking genome-scale metabolic modeling and genome annotation. Methods Mol Biol 985, 61–83.

Bukau, B., and Horwich, A.L. (1998). The Hsp70 and Hsp60 chaperone machines. Cell 92, 351–366.

Chapman, E., Farr, G.W., Usaite, R., Furtak, K., Fenton, W.A., Chaudhuri, T.K., Hondorp, E.R., Matthews, R.G., Wolf, S.G., Yates, J.R., et al. (2006). Global aggregation of newly translated proteins in an Escherichia coli strain deficient of the chaperonin GroEL. Proc Natl Acad Sci U S A 103, 15800–15805.

Chilukoti, N., Kumar, C.M., and Mande, S.C. (2015). GroEL2 of Mycobacterium tuberculosis Reveals the Importance of Structural Pliability in Chaperonin Function. J Bacteriol 198, 486–497.

Durao, P., Aigner, H., Nagy, P., Mueller-Cajar, O., Hartl, F.U., and Hayer-Hartl, M. (2015). Opposing effects of folding and assembly chaperones on evolvability of Rubisco. Nat Chem Biol 11, 148–155.

Faith, J.J., Driscoll, M.E., Fusaro, V.A., Cosgrove, E.J., Hayete, B., Juhn, F.S., Schneider, S.J., and Gardner, T.S. (2008). Many Microbe Microarrays Database: uniformly normalized Affymetrix compendia with structured experimental metadata. Nucleic Acids Res 36, D866–870.

Fares, M.A., Ruiz-Gonzalez, M.X., Moya, A., Elena, S.F., and Barrio, E. (2002). Endosymbiotic bacteria: groEL buffers against deleterious mutations. Nature 417, 398.

Fayet, O., Ziegelhoffer, T., and Georgopoulos, C. (1989). The groES and groEL heat shock gene products of Escherichia coli are essential for bacterial growth at all temperatures. J Bacteriol 171, 1379–1385.

Ferrer, M., Chernikova, T.N., Yakimov, M.M., Golyshin, P.N., and Timmis, K.N. (2003). Chaperonins govern growth of Escherichia coli at low temperatures. Nat Biotechnol 21, 1266–1267.

Guisbert, E., Herman, C., Lu, C.Z., and Gross, C.A. (2004). A chaperone network controls the heat shock response in E. coli. Genes Dev 18, 2812–2821.

Guzman, L.M., Belin, D., Carson, M.J., and Beckwith, J. (1995). Tight regulation, modulation, and high-level expression by vectors containing the arabinose PBAD promoter. J Bacteriol 177, 4121–4130.

Hartl, F.U., and Hayer-Hartl, M. (2002). Molecular chaperones in the cytosol: from nascent chain to folded protein. Science 295, 1852–1858.

Hirosawa, M., Hoshida, M., Ishikawa, M., and Toya, T. (1993). MASCOT: multiple alignment system for protein sequences based on three-way dynamic programming. Comput Appl Biosci 9, 161–167.

Horwich, A.L., Low, K.B., Fenton, W.A., Hirshfield, I.N., and Furtak, K. (1993). Folding in vivo of bacterial cytoplasmic proteins: role of GroEL. Cell 74, 909–917.

Houry, W.A., Frishman, D., Eckerskorn, C., Lottspeich, F., and Hartl, F.U. (1999). Identification of in vivo substrates of the chaperonin GroEL. Nature 402, 147–154.

Ishimoto, T., Fujiwara, K., Niwa, T., and Taguchi, H. (2014). Conversion of a chaperonin GroEL-independent protein into an obligate substrate. J Biol Chem 289, 32073–32080.

Kandror, O., Sherman, M., Moerschell, R., and Goldberg, A.L. (1997). Trigger factor associates with GroEL in vivo and promotes its binding to certain polypeptides. J Biol Chem 272, 1730–1734.

Kandror, O., Sherman, M., Rhode, M., and Goldberg, A.L. (1995). Trigger factor is involved in GroEL-dependent protein degradation in Escherichia coli and promotes binding of GroEL to unfolded proteins. EMBO J 14, 6021–6027.

Kayser, A., Weber, J., Hecht, V., and Rinas, U. (2005). Metabolic flux analysis of Escherichia coli in glucose-limited continuous culture. I. Growth-rate-dependent metabolic efficiency at steady state. Microbiology 151, 693–706.

Kerner, M.J., Naylor, D.J., Ishihama, Y., Maier, T., Chang, H.C., Stines, A.P., Georgopoulos, C., Frishman, D., Hayer-Hartl, M., Mann, M., et al. (2005). Proteome-wide analysis of chaperonin-dependent protein folding in Escherichia coli. Cell 122, 209–220.

Kim, Y.E., Hipp, M.S., Bracher, A., Hayer-Hartl, M., and Hartl, F.U. (2013). Molecular chaperone functions in protein folding and proteostasis. Annu Rev Biochem 82, 323–355.

Kumar, C.M., Mande, S.C., and Mahajan, G. (2015). Multiple chaperonins in bacteria--novel functions and non-canonical behaviors. Cell Stress Chaperones 20, 555–574.

Kumar, C.M.S. (2017). Prokaryotic Multiple Chaperonins: The Mediators of Functional and Evolutionary Diversity. In Prokaryotic Chaperonins: Multiple Copies and Multitude Functions, C.M.S. Kumar, and S.C. Mande, eds. (Singapore: Springer Singapore), pp. 39–51.

Kusukawa, N., and Yura, T. (1988). Heat shock protein GroE of Escherichia coli: key protective roles against thermal stress. Genes Dev 2, 874–882.

Lemos, J.A., Luzardo, Y., and Burne, R.A. (2007). Physiologic effects of forced down-regulation of dnaK and groEL expression in Streptococcus mutans. J Bacteriol 189, 1582–1588.

Li, G.W., Burkhardt, D., Gross, C., and Weissman, J.S. (2014). Quantifying absolute protein synthesis rates reveals principles underlying allocation of cellular resources. Cell 157, 624–635.

Llorca, O., Galan, A., Carrascosa, J.L., Muga, A., and Valpuesta, J.M. (1998). GroEL under heat-shock. Switching from a folding to a storing function. J Biol Chem 273, 32587–32594.

Lund, P.A. (2001). Molecular Chaperones in the Cell (Oxford University Press).

Lund, P.A. (2009). Multiple chaperonins in bacteria--why so many? FEMS Microbiol Rev 33, 785–800.

Macho, A.P., Guidot, A., Barberis, P., Beuzon, C.R., and Genin, S. (2010). A competitive index assay identifies several Ralstonia solanacearum type III effector mutant strains with reduced fitness in host plants. Mol Plant Microbe Interact 23, 1197–1205.

Martin, J., Horwich, A.L., and Hartl, F.U. (1992). Prevention of protein denaturation under heat stress by the chaperonin Hsp60. Science 258, 995–998.

Martinez-Alonso, M., Garcia-Fruitos, E., Ferrer-Miralles, N., Rinas, U., and Villaverde, A. (2010). Side effects of chaperone gene co-expression in recombinant protein production. Microb Cell Fact 9, 64.

Monk, I.R., Casey, P.G., Cronin, M., Gahan, C.G., and Hill, C. (2008). Development of multiple strain competitive index assays for Listeria monocytogenes using pIMC; a new site-specific integrative vector. BMC Microbiol 8, 96.

Musatovova, O., Dhandayuthapani, S., and Baseman, J.B. (2006). Transcriptional heat shock response in the smallest known self-replicating cell, Mycoplasma genitalium. J Bacteriol 188, 2845–2855.

O’Leary, N.A., Wright, M.W., Brister, J.R., Ciufo, S., Haddad, D., McVeigh, R., Rajput, B., Robbertse, B., Smith-White, B., Ako-Adjei, D., et al. (2016). Reference sequence (RefSeq) database at NCBI: current status, taxonomic expansion, and functional annotation. Nucleic Acids Res 44, D733–745.

Orth, J.D., Conrad, T.M., Na, J., Lerman, J.A., Nam, H., Feist, A.M., and Palsson, B.O. (2011). A comprehensive genome-scale reconstruction of Escherichia coli metabolism--2011. Mol Syst Biol 7, 535.

Sabater-Munoz, B., Prats-Escriche, M., Montagud-Martinez, R., Lopez-Cerdan, A., Toft, C., Aguilar-Rodriguez, J., Wagner, A., and Fares, M.A. (2015). Fitness Trade-Offs Determine the Role of the Molecular Chaperonin GroEL in Buffering Mutations. Mol Biol Evol.

Samuelson, J.C. (2011). Recent developments in difficult protein expression: a guide to E. coli strains, promoters, and relevant host mutations. Methods Mol Biol 705, 195–209.

Schumann, W. (2017). Regulation of the Heat Shock Response in Bacteria. In Prokaryotic Chaperonins: Multiple Copies and Multitude Functions, C.M.S. Kumar, and S.C. Mande, eds. (Singapore: Springer Singapore), pp. 21–36.

Singhal, K., Vreede, J., Mashaghi, A., Tans, S.J., and Bolhuis, P.G. (2015). The Trigger Factor Chaperone Encapsulates and Stabilizes Partial Folds of Substrate Proteins. PLoS Comput Biol 11, e1004444.

Smith, H.L. (2011). Bacterial competition in serial transfer culture. Math Biosci 229, 149–159.

Tokuriki, N., Stricher, F., Serrano, L., and Tawfik, D.S. (2008). How protein stability and new functions trade off. PLoS Comput Biol 4, e1000002.

Tokuriki, N., and Tawfik, D.S. (2009). Chaperonin overexpression promotes genetic variation and enzyme evolution. Nature 459, 668–673.

UniProt, C. (2019). UniProt: a worldwide hub of protein knowledge. Nucleic Acids Res 47, D506–D515.

Van Dyk, T.K., Gatenby, A.A., and LaRossa, R.A. (1989). Demonstration by genetic suppression of interaction of GroE products with many proteins. Nature 342, 451–453.

van Opijnen, T., and Camilli, A. (2013). Transposon insertion sequencing: a new tool for systems-level analysis of microorganisms. Nat Rev Microbiol 11, 435–442.

Vulic, M., and Kolter, R. (2001). Evolutionary cheating in Escherichia coli stationary phase cultures. Genetics 158, 519–526.

Wang, M., Weiss, M., Simonovic, M., Haertinger, G., Schrimpf, S.P., Hengartner, M.O., and von Mering, C. (2012). PaxDb, a database of protein abundance averages across all three domains of life. Mol Cell Proteomics 11, 492–500.

Weber, J., Kayser, A., and Rinas, U. (2005). Metabolic flux analysis of Escherichia coli in glucose-limited continuous culture. II. Dynamic response to famine and feast, activation of the methylglyoxal pathway and oscillatory behaviour. Microbiology 151, 707–716.

Weiner, J., 3rd, Zimmerman, C.U., Gohlmann, H.W., and Herrmann, R. (2003). Transcription profiles of the bacterium Mycoplasma pneumoniae grown at different temperatures. Nucleic Acids Res 31, 6306–6320.

Williams, T.A., and Fares, M.A. (2010). The effect of chaperonin buffering on protein evolution. Genome Biol Evol 2, 609–619.

Wyganowski, K.T., Kaltenbach, M., and Tokuriki, N. (2013). GroEL/ES buffering and compensatory mutations promote protein evolution by stabilizing folding intermediates. J Mol Biol 425, 3403–3414.

Zambrano, M.M., Siegele, D.A., Almiron, M., Tormo, A., and Kolter, R. (1993). Microbial competition: Escherichia coli mutants that take over stationary phase cultures. Science 259, 1757–1760.

Zou, T., Williams, N., Ozkan, S.B., and Ghosh, K. (2014). Proteome folding kinetics is limited by protein halflife. PLoS One 9, e112701.

