## Supplementary material for "Chaperonin Abundance Boosts Bacterial Fitness": File 1

* Address Correspondence to

**Running Title:** Chaperonin Enhances Fitness.

**Key words:** Metabolic flux, Chaperonin, GroEL, Evolution, Proteomics, Metabolism

### Supplementary Methods

**Growth Parameters of GL-$L_b$ and GL-$H_b$ Strains.** Overnight cultures of GL-$L_b$ and GL-$H_b$ strains were inoculated at equal cell number into two flasks with fresh LB supplemented with 0.2% L-arabinose and 0.2% D-lactose and were incubated with constant shaking at 30 °C. $OD_{600}$ was measured for both the cultures at constant intervals and logarithmic values for the cell numbers at each interval were calculated. The growth parameters, such as growth rate, doubling time and growth rate constant, were calculated as per the standard calculations (Neidhardt et al., 1990). Briefly, growth rate constant was calculated as the rate of difference in the number of cells between the initial and final stages of exponential growth, using the formula, $\mu = (\log10\ N - \log10\ N0) \times 2.303) / (t - t0)$. Doubling time (g) was calculated using the formula - $g = (0.693/\mu) \times 60$. Growth rate (k) was calculated by dividing μ by 0.693.

**Flux Balance Analysis of the GL-$L_b$ and GL-$H_b$ Strains to assess metabolic changes.** The *E. coli* genome-scale metabolic network iJO1366 (Orth et al., 2011) was used for the Flux Balance Analysis (FBA) simulations. Here, a simple linear relationship between the enzyme abundance (X) and the flux (F) through a corresponding reaction was assumed. TransFlux, a software tool developed in-house (available at: http://www.nccs.res.in/TransFlux/index.jsp) was used to overlay the enzyme abundance on the FBA simulations.

The iJO1366 model, the first simulated FBA model, employed standard energy source (equivalent to a glucose-supplemented minimal media) to obtain the steady state fluxes through all the reactions (Orth et al., 2011). The objective function of the FBA simulation was to maximize the biomass production, while using some 'default constraints' (lower- and upper-bounds of fluxes through each reaction) derived from the literature (Blais et al., 2013). The fluxes obtained for each of the reactions were considered to constitute the set of 'reference fluxes' ($F_R$). Subsequently, for a subset of reactions, the upper-bounds (UB) and lower-bounds (LB) of the fluxes (i.e. the reaction constraints) were re-computed based on the putative expression levels of the corresponding enzyme-encoding genes (Supplementary File 2). In addition to the results from the current proteomic study, *E. coli* gene expressions, derived from the Many Microbe Microarrays Database (M3D, www.m3d.mssm.edu) (Faith et al. 2008) were screened, to gauge the possible extent of fluctuations in the expression levels of the enriched metabolic enzymes, in GL-$L_b$ and GL-$H_b$ strains. The ratio between the maximum and minimum expression values for specific enzymes, as obtained from the M3D, were considered as fold change (FC) and used to derive differential fluxes through corresponding reactions of the GL-$L_b$ and GL-$H_b$ strains. It may be noted that spots for some of the enzymes were not identified in the 2D-PAGE analysis in either GL-$L_b$ or GL-$H_b$ strains, although they were observed to have significant expression levels in the micro-array data. For these cases, the 'down-regulated' expression value of the enzyme was considered to be 0 (i.e. equivalent to no expression/ a deletion mutant). While re-computing the reaction constraints, it was assumed that the different expression levels of the enzyme X (say $X_1$ and $X_2$ respectively), corresponding to GL-$L_b$ (say condition 1) and GL-$H_b$ (say condition 2) strains, had a linear relationship with the corresponding fluxes through the reaction ($F_1$ and $F_2$, respectively). Consequently, the fold change ($FC_{1:2}$) in expression values of $X_1$ and $X_2$, would also correspond to the fold change in reaction fluxes, as shown below.

$$\frac{X_1}{X_2} = FC_{1:2} = \frac{F_1}{F_2} \ldots\ldots\ldots\ldots\ldots\ldots\ldots. (i)$$

Therefore, $F_1$ and $F_2$ may be expressed in terms of $FC_{1:2}$ as, $F_1 = F_2 * FC_{1:2}$ ……………………..(ii) $F_2 = \frac{F_1}{FC_{1:2}}$ ……………………..(iii)

It was further assumed that the reference flux ($F_R$) through a reaction corresponded to a condition where a given enzyme (X) was neither 'upregulated' nor 'downregulated', significantly. $F_R$ may therefore be considered to have an intermediate value, between $F_1$ and $F_2$, and was assumed to be a simple average of $F_1$ and $F_2$, as shown below -

$$F_R = \frac{F_1 + F_2}{2} \dots\dots\dots\dots\dots\dots\dots\dots (iv)$$

From equations ii, iii and iv, the values of $F_1$ and $F_2$ may be calculated as,

$$F_1 = \frac{2 * F_R}{\left(1 + \frac{1}{FC_{1:2}}\right)} \dots\dots\dots\dots\dots\dots\dots\dots (v)$$

$$F_2 = \frac{2 * F_R}{(1 + FC_{1:2})} \dots\dots\dots\dots\dots\dots\dots\dots (vi)$$

During the conditions 1 and 2 FBA simulations, the UB and LB of the fluxes through relevant reactions were constrained based on the values of $F_i$ (i.e., $F_1$ or $F_2$, depending on the chosen condition) and the reference flux $F_R$ - if, $F_i > F_R$, then the LB was constrained to $F_i$ and the UB was left unchanged; else if, $F_i < F_R$, then the UB was constrained to $F_i$ and the LB was left unchanged.

For multi-enzyme catalyzed reactions, the effect of 'constrained bounds' were compounded using logical operators, such as 'AND' and 'OR'. While the AND operator was used to represent a scenario, wherein all the multiple enzymes were necessary to catalyze a single reaction, the OR operator represented reactions catalyzed by orthologous enzymes. Thus, a cumulative value pertaining to expression of the enzymes involved in such reactions was calculated as shown below. This value was subsequently used for defining the LB/UB in accordance with equations described above.

$X^{ab} = min(X^a, X^b)$ … … … when enzymes ‘a’ AND ‘b’ both are essential for the reaction

$X^{ab} = max(X^a, X^b)$ … … … when either of enzymes ‘a’ OR ‘b’ is required for the reaction

Furthermore, while performing two different FBA simulations, for conditions 1 and 2, the biomass production levels of the strains were constrained into a narrow range of ± 25% of their experimentally observed values (Fig. S4).

***In silico* Identification of Proteins Co-expressed with GroE.** Publicly available proteomic data corresponding to *E. coli* was collated from the paxdb database (http://pax-db.org/) and screened for proteins potentially co-enriched with GroEL across different experiments, as below. Pearson correlation values between the expression levels of GroEL and all other protein-encoding genes were computed. Protein-encoding genes exhibiting significantly correlated expression with GroE (Pearson correlation co-efficient >= 0.7, $P < 0.05$) were selected for further analyses. Affiliations of these proteins to different COG classes were ascertained and cumulative statistics for each of the COG categories were computed (poorly characterized proteins were not considered). Moreover, Gene Ontology (GO) enrichment analysis of this set of proteins were performed with the DAVID tool (https://david.ncifcrf.gov/), to ascertain their repertoire of biological processes. GO biological process terms (level 3) satisfying a count of more than ten genes and an ‘*ease*’ value of 0.1, were considered.

**Assessment of Response to Lowered Aeration.** Stationary phase cultures of *E. coli* strains, GL-$L_b$ and GL-$H_b$, were normalized and sub-cultured at equal cell number into two sets of LB broth supplemented as above. One set was incubated with constant shaking at 200 x RPM and the other set was overlaid with mineral oil and incubated as standing culture. These two sets, representing optimally and minimally aerated cultures, respectively, were incubated at 30 °C overnight. To measure the extent of growth in the optimally aerated and low aerated GL-$L_b$ and GL-$H_b$ cultures, the resulting cultures were serially diluted in LB and were spotted onto LB agar plates supplemented with 0.2% D-lactose, to induce the chromosomal *groE* operon and allow the growth of both GL-$L_b$ and GL-$H_b$ strains. The plates were incubated at 30 °C.

### Supplementary Results

**Flux Balance Analysis reveals a basis for Cellular Fitness of the GroE Overproducing Strain.** Considering the preferential enrichment of metabolic enzymes upon GroE overproduction, we adopted an FBA approach (Blais et al., 2013; Orth et al., 2011) to assess how the differential enrichment of metabolic enzymes in GL-$L_b$ and GL-$H_b$ strains would translate into altered metabolic states and cellular fitness (Supplementary File 1). A genome-scale metabolic model of *E. coli*, viz. iJO1366 (Orth et al., 2011), was used for the FBA simulations. During the simulations, it was assumed that an abundant enzyme will proportionally enrich the metabolic flux through a corresponding reaction. Since flux through any given metabolic reaction is dependent on the intracellular abundance of the catalyzing enzyme, gene expression and/or proteome data could be utilized, in principle, to constrain reaction fluxes in a genome-scale metabolic model. Therefore, in addition to the results from the current proteomic study, observations from *E. coli* gene expression microarray studies, derived from M3D (Faith et al., 2008), were screened to gauge the extent of fluctuations in the expression levels of the enriched metabolic enzymes, in the two *E. coli* strains. This information was subsequently utilized to constrain fluxes through respective reactions while performing two independent FBA simulations, each of which corresponded to the enzyme expression/enrichment profiles of GL-$L_b$ and GL-$H_b$ strains. The enzyme levels were incorporated into FBA simulations using in-house developed tool, 'TransFlux' (http://www.nccs.res.in/TransFlux/index.jsp).

Average flux through the reactions constituting different metabolic pathways were obtained through the FBA simulations on the GL-$L_b$ and GL-$H_b$ strains (Table 2). As expected, higher flux was observed in the simulated GL-$H_b$ strain through several pathways of carbon metabolism including glycolysis, gluconeogenesis, citric acid cycle (TCA cycle) and its anaplerotic reactions, as well as alternate carbon metabolism (Table S2). Apparently, flux through these pathways was enhanced by elevated import of glucose and glycerol (Supplementary File 2). Moreover, pathways corresponding to the metabolism of several glucogenic amino acids and energy generating oxidative phosphorylation were all enriched in this strain. Interestingly, the pathways leading to the metabolism of methylglyoxal, a toxic side product of several metabolic pathways, including glycolysis, were enriched in the GL-$H_b$ strain (Table S2). However, pathways leading to the metabolism of membrane lipids, pyruvic acid, pentose sugars, ubiquinone and salvage of nucleotides were enriched in the GL-$L_b$ strain.

The enhanced reactions in the said pathways exhibited a correlation between the pathways and growth. Enhancement of the reactions of the methylglyoxal biosynthetic pathway that are known to be triggered when the cells have abundant nutrients (Kayser et al., 2005; Weber et al., 2005), further confirms the elevated energy status of the GL-$H_b$ cells. Some of the side reactions from the amino acid metabolism, such as that catalyzed by L-threonine dehydrogenase and the follow-up reaction catalyzed by Aminoacetone:oxygen oxidoreductase

(FMN), appeared to be enriched in the GL-$H_b$ strain (Supplementary File 2). Moreover, increased excretion of L-lactic acid in combination with an enhanced glycolytic side reaction, catalyzed by L-Lactaldehyde:$NADP^+$ 1-oxidoreductase, indicated a probable increase in the intracellular methylglyoxal concentrations. Interestingly, increased excretion of the fermentation products, such as ethanol and acetic acid (Supplementary File 2), suggested the presence of an acetate switch (Wolfe, 2005) in this strain. Although in these simulations formic acid is excreted from both the strains, excretion from GL-Lb strain was higher.

In addition, the enrichment of L-glutamate metabolism, especially by the reaction catalyzed by glutamate dehydrogenase, the branch-point enzyme between carbon and nitrogen metabolisms catalyzing the anaplerotic reaction in the aminating direction, i. e., towards the biosynthesis of Glutamine, connected the pathways leading to the biosynthesis of the other glucogenic amino acids (Supplementary File 2). Enhanced levels of oxygen-uptake and probable increased $CO_2$ expulsion further supported the observed enhancement in oxidative phosphorylation reactions. These observations suggested an increased respiration in GL-$H_b$ relative to GL-Lb

However, some of the reactions enhanced in the GL-$L_b$ strain appear to compensate for some metabolic deficiencies in this strain. The reactions that lead to breakdown of long-chain fatty acids, such palmetoyl-CoA, stearoyl-CoA and 3-hydroxystearoyl-CoA, which are synthesized by the reactions of acetyl-CoA acyltransferase, acyl-CoA dehydrogenase and 3-hydroxyacyl-CoA dehydratase, respectively, were enriched in this strain. Moreover, the reactions leading to the salvage of nucleotides from the degradation intermediates and their feed reactions, such as ubiquinone derivative biosynthesis, pentose phosphate pathway and the reactions catalyzed by

NDP kinase and NDP reductase, appeared to be enriched in this strain (Supplementary File 2). In addition, several reactions of the pyruvate metabolism that act as alternate feed reactions for TCA cycle and amino acid metabolism, were enriched. For example, the reactions catalyzed by pyruvate synthase and acetyl-CoA synthetase, that act as alternate feed reactions for L-alanine/L-asparagine biosynthesis and TCA cycle, respectively were enriched. Overall, FBA simulations (Supplementary File 2) indicated an enriched carbon, nitrogen and energy metabolism in the GL-$H_b$ strain, suggesting an enhancement in energy metabolism due to chaperonin overproduction and further strengthening our perception of a direct relation between chaperonins and cellular fitness.

### Supplementary Discussion

The FBA revealed a correlation between chaperonin abundance and cellular metabolism. Enhanced import of metabolites and precursors into GL-$H_b$ cells indicate that the strain is able to conserve energy that otherwise would be utilized in synthesizing these molecules. Moreover, FBA could predict the enhancement of nucleotide salvage pathway and its feeder pathways in GL-$L_b$ cells (Table S2 and Supplementary File 2). Enhanced carbon and nitrogen metabolism in GL-$H_b$ strain, accompanied with the biosynthesis of several glucogenic amino acids is predicted lead to an increase in available energy and consequent enhanced growth or stress resistance in this strain, consistent with the experimental data (Table S2 and Supplementary File 2). The GL-$L_b$ cells apparently rely on energetically less favorable alternative pathways such as pyruvate metabolism, pentose phosphate pathway, and nucleotide salvage pathways that would feed into the major metabolic pathways in synthesizing these metabolites.

In addition, operation of the acetate switch and the consequent increased excretion of the fermentation products, such as acetic and formic acids from the GL-$H_b$ and GL-$L_b$ strains (Supplementary File 2), respectively, indicate a strong correlation of metabolic and oxidative status. Accumulation and excretion of acetic acid has been demonstrated in nutrient rich and fast growing cultures. It is triggered by excess nutrients and consequent accumulation of acetyl CoA (Wolfe, 2005). In addition to carbohydrate metabolism, enhanced flux through reaction catalyzed by acetyl CoA ligase in the GL-$H_b$ strain (Supplementary File 2) supports the hypothesis that GL-$H_b$ cells might be nutrient-rich. However, increased formate accumulation has been demonstrated to increase the activity of pyruvate formatelyase (PFL) in pyruvate metabolism that converts acetyl-CoA to pyruvate (Alexeeva et al., 2000) and understandably, depletion of this central molecule of metabolism will repress several metabolic pathways, such as the TCA cycle. Notably, the said features - increased excretion of formate, increased activity of pyruvate formatelyase and depletion in TCA cycle - are apparent in the simulated GL-$L_b$ strain (Supplementary File 2). Enhancement of the PFL reaction (Supplementary File 2) that is known to be enhanced during anaerobic growth (de Graef et al., 1999), also predicts diminished growth of GL-$L_b$ under oxygen-limiting conditions, as is seen experimentally (Fig. S2). Interestingly, the differentially enriched proteins (Table S1) correlate with the acetate switch (Supplementary File 2). Maltose-binding periplasmic protein and OmpC are known to be upregulated during formate and acetate stress, respectively (Kirkpatrick et al., 2001), as observed here. Taken together, these results show how the growth and redox phenotypes of the GL-$H_b$ and GL-$L_b$ strains seen experimentally may arise from their respective metabolic condition as modelled in FBA from the proteomic data.

Moreover, confirming the notion that GroEL-GroES depletion results in defective membrane biogenesis (Fujiwara and Taguchi, 2007; McLennan and Masters, 1998), several reactions in membrane lipid metabolism that result in the breakdown of long chain fatty acids are enriched in the GL-$L_b$ strain, which ultimately leads to increased pools of acetyl-CoA that can further feed to several important metabolic pathways. This also indicates that the GL-$L_b$ strain is compromised for several metabolic pathways and thus is less robust than the GL-$H_b$ strain. FBA indicates a greater uptake of oxygen and higher flux through $CO_2$ releasing reactions in the GL-$H_b$ cells, and therefore suggest a more optimal state of respiration (Table 2 and Supplementary File 2). The dilution spotting assay (Fig. S4) showed that the chaperonin deficient cells grow less well under lower aeration. Taken together, these studies provide direct rationales for the observed effect of chaperonins on the cellular proteome and fitness.

### Supplementary Figures and Legends

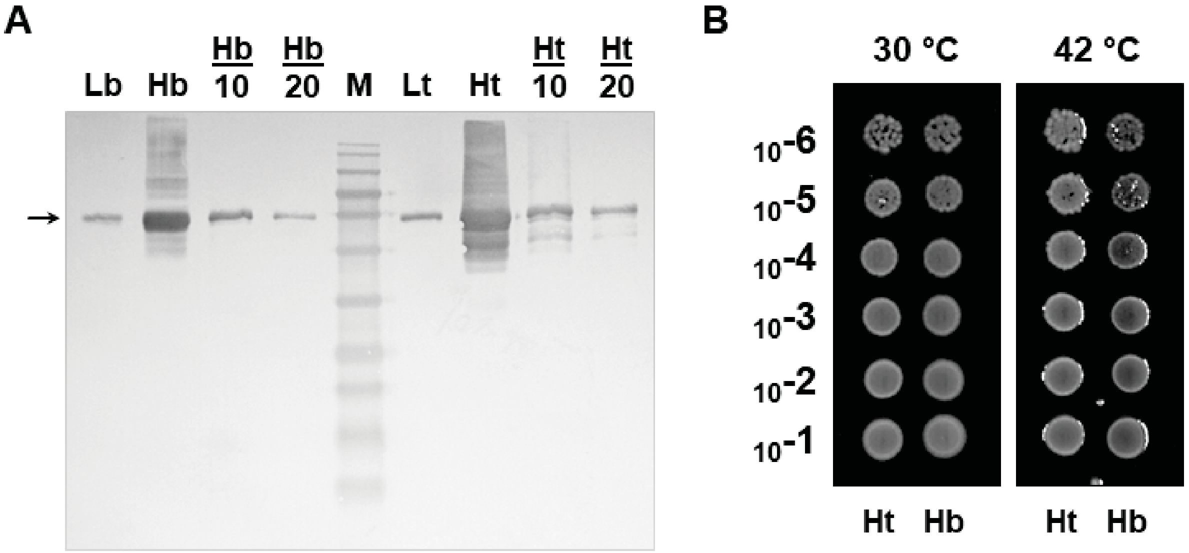


**Figure S1. The pTrc99a and pBAD24 based strains show comparable characteristics. A.** Enhanced GroEL production in GL-$H_t$ and GL-$H_b$ compared to their vector-only strains. Soluble lysates of GL-$L_b$ (Lb), GL-$H_b$ (Hb), GL-$H_t$ (Ht) and GL-$L_t$ (Lt) were resolved on a 12.5% SDS-PAGE and probed with GroEL specific antibody. Hb/10, Hb/20, Ht/10 and Ht/20 indicate the lanes with ten and twenty fold diluted lysates of GL-$H_b$ and GL-$H_t$, respectively. **B.** GL-$H_t$ and GL-$H_b$ strains show similar temperature resistance. Serially diluted cultures of the GL-$H_t$ (Ht) and GL-$H_b$ (Hb) strains were spotted on to the LB agar plates and incubated at the indicated temperatures.

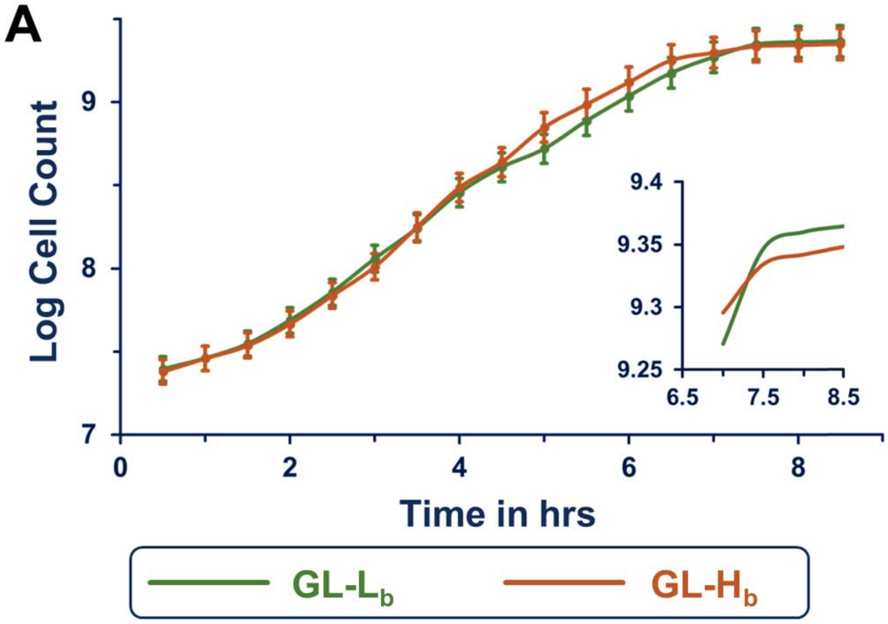


**B**

| Strain | Growth rate constant (μ) | Growth rate (hr $^{-1}$) | Doubling time (min) |
|---|---|---|---|
| GL-$L_b$ | 1.62 ± 0.12 | 2.33 ± 0.17 | 25:52 ± 1.94 |
| GL-$H_b$ | 1.81 ± 0.08 | 2.60 ± 0.12 | 23:07 ± 1.08 |

**Figure S2. GL-$H_b$ and GL-$L_b$ Strains Exhibit Comparable Growth Parameters. A.** Growth curves depicting an increase in the population of the indicated strains as a function of time. The strains were cultured at 30 °C in standard LB supplemented as appropriate. The curves were fit using the least-squared method. The sub-plot with a zoomed-up region of the growth curve depicts the stationary phase of the indicated cultures. **B.** Table showing the principal growth parameters of the indicated cultures. The values are average of four independent experiments.

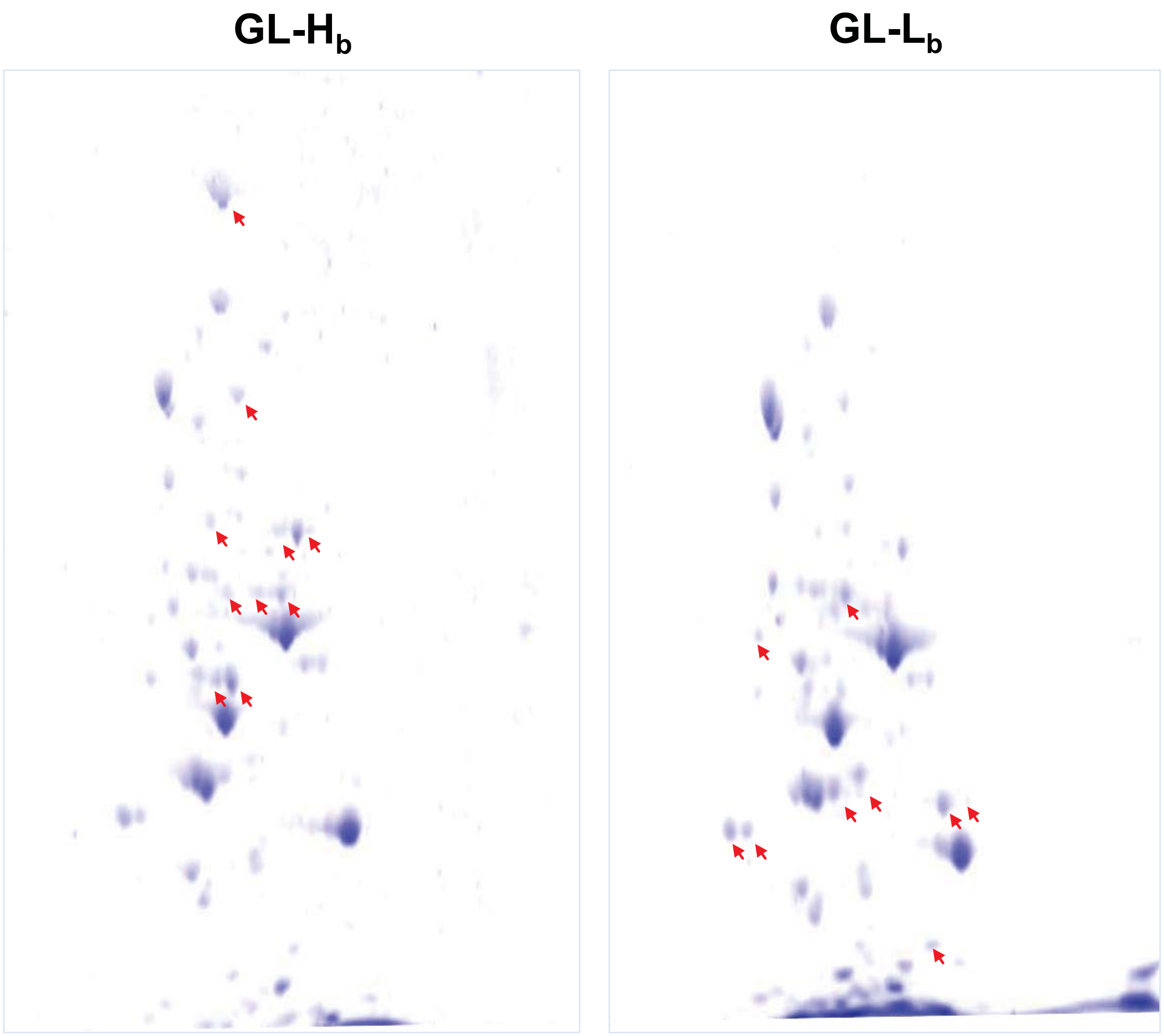


**Figure S3. 2D PAGE analysis of the soluble proteomes of GL-H$_b$ and GL-L$_b$ Strains. Soluble lysates** of **GL-H$_b$ and GL-L$_b$ Strains** were resolved on a 2D PAGE, with the first and second dimensions through a 3 - 10 pH gradient strip and 10% SDS-PAGE, respectively. The separated proteins were stained with Coomassie brilliant blue and intensities of the stained protein spots were compared between the two gels using densitometry. Arrows indicate differentially enriched spots.

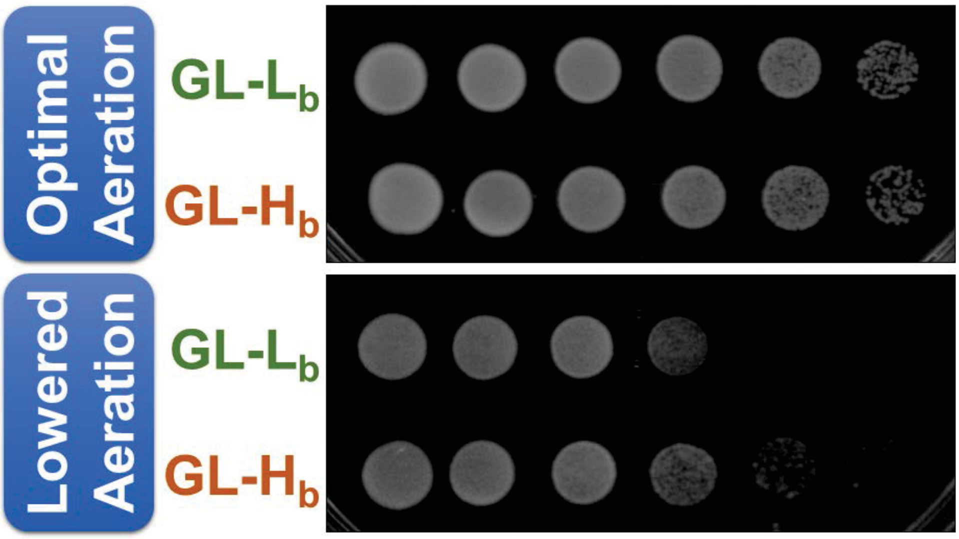


**Figure S4. GL-$L_b$, but not GL-$H_b$, showed lower growth under low oxygen conditions.** To evaluate the effect of aeration on the growth of the indicated strains, shaking and standing oil-overlaid liquid cultures, which represent optimal and minimal aeration, respectively, were incubated at 30 °C overnight. To measure the bacterial growth in the indicated conditions, the resultant cultures were serially diluted and spotted on LB agar plates, supplemented with lactose.

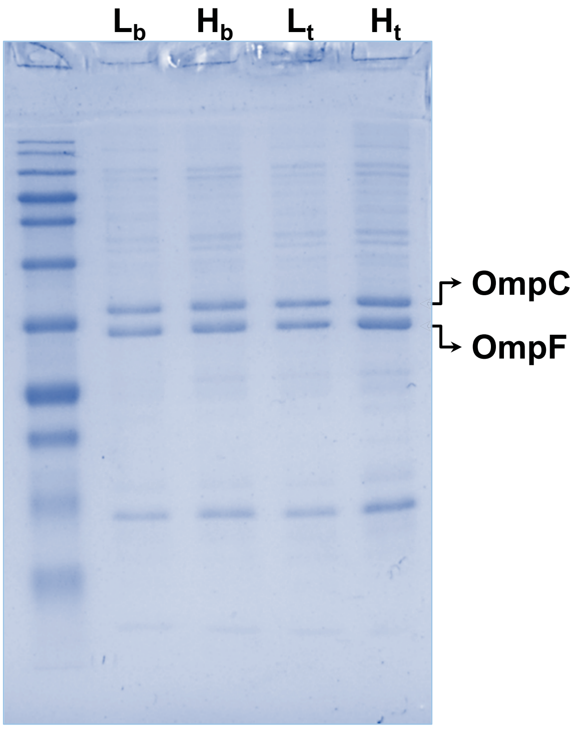


**Figure S5. Cell Fractionation Revealed Differential Levels of outer membrane proteins.** Cells of GL-$H_b$ ($H_b$), GL-$L_b$ ($L_b$), GL-$H_t$ ($H_t$) and GL-$L_t$ ($L_t$) strains were fractionated by high-speed centrifugation following sonication to isolate membrane fractions. These fractions were resolved on 12.5% SDS-PAGE to reveal membrane protein profiles of the indicated strains.

**Table S1. Properties of the Differentially Enriched Proteins in GL-Hb and GL-Lb Strains.**

| Strain | SwissProt Entry Name | SwissProt Accession Number | Protein Description | MW [kDa] | pI | Unique Peptides | Coverage | GroEL Substrate Class* | Oligomeric State | COG Functional Category | SCOP Fold | Location *in vivo* (SwissProt) | Gene names | Essentiality | mRNA $t_{1/2}$ (min) |
|---|---|---|---|---|---|---|---|---|---|---|---|---|---|---|---|
| GL-$H_b$ | ENO_ECOLI | P0A6P9 | Enolase (EC:4.2.1.11) (2-phosphoglycerate dehydratase) (2-phospho-D-glycerate hydro-lyase) | 45.5 | 5.32 | 19 | 51.68 | One | Homodimer | G | c.1.1.1; d.54.1.1 | Cytoplasmic, cytoskeleton, secreted, cell surface | *eno (b2779)* | 1 | 4.7 |
| | 6PGD_ECOLI | P00350 | 6-phosphogluconate dehydrogenase, decarboxylating (EC:1.1.1.44) | 51.5 | 5.04 | 16 | 57.48 | One | Homodimer | G | a.100.1; c.2.1.6 | Cytoplasm | *gnd (b2029)* | 0 | 10.6 |
| | DLDH_ECOLI | P0A9P0 | Dihydrolipoyl dehydrogenase (EC:1.8.1.4), Dihydrolipoamide dehydrogenase, E3 component of pyruvate and 2-oxoglutarate dehydrogenases complexes | 50.6 | 5.79 | 18 | 44.51 | Four | Homodimer | C | d.87.1.1; c.3.1.5 | Cytoplasmic, cell inner membrane, peripheral membrane protein | *lpdA (b0116)* | 1 | 5.8 |
| | MDH_ECOLI | P61889 | Malate dehydrogenase (EC:1.1.1.37) | 32.3 | 5.28 | 19 | 87.5 | Two | Homodimer | C | d.162.1.1; c.2.1.5 | Cytoplasmic | *mdh (b3236)* | 1 | 10.5 |
| | TYPH_ECOLI | P07650 | Thymidine phosphorylase (EC:2.4.2.4) | 51.4 | 5.2 | 19 | 51.36 | Four | Homodimer | F | c.27.1.1; d.41.3.1; a.46.2.1 | Cytoplasmic | *deoA (b4382)* | 0 | 15.8 |
| | IDH_ECOLI | P08200 | Isocitrate dehydrogenase [NADP] (EC:1.1.1.42) (Oxalosuccinate decarboxylase) | 45.7 | 5.15 | 19 | 65.28 | - | Homodimer | C | c.77.1.1 | Cytoplasmic | *icd (b1136)* | 0 | 5.8 |
| | ACEA_ECOLI | P0A9G6 | Isocitrate lyase (EC:4.1.3.1) | 47.4 | 5.16 | 11 | 40.32 | - | Homotetramer | C | c.1.12.7 | Cytoplasmic | *aceA (b4015)* | 0 | 11.5 |
| | DPO3B_ECOLI | P0A988 | DNA polymerase III beta subunit protein (EC:2.7.7.7) | 40.5 | 5.45 | 12 | 40.71 | - | Hetero-Oligomer | L | d.131.1.1 | Cytoplasmic | *dnaN (b3701)* | 1 | 2.4 |
| | TALB_ECOLI | P0A870 | Transaldolase B (EC:2.2.1.2) | 35 | 5.11 | 21 | 77.29 | Five | Homodimer | G | c.1.10.1 | Cytoplasmic | *talB (b0008)* | 0 | 3.4 |
| | POTD_ECOLI | P0AFK9 | Spermidine/putrescine-binding periplasmic protein | 38.8 | 4.86 | 16 | 50.86 | - | Monomer | E | c.94.1.1 | Periplasmic | *potD (b1123)* | 1 | - |
| | RIHA_ECOLI | P41409 | Pyrimidine-specific ribonucleoside hydrolase, RihA (EC:3.2.2.-), Cytidine/uridine-specific hydrolase, ribonucleoside hydrolase 1 | 33.8 | 4.84 | 13 | 77.81 | - | Tetramer | F | C.70.1.0 | Cytoplasmic | *rihA (b0651)* | 0 | 4.4 |
| | CH60_ECOLI | P0A6F5 | Chaperonin 60, GroEL | 57.0 | 4.85 | 11 | 28.89 | Five | Homo-tetradecamer | O | a.129.1.1; d.56.1.1; c.8.5.1 | Cytoplasmic | *groL (b4143)* | 1 | 3.5 |
| | BGAL_ECOLI | P00722 | Beta-galactosidase (EC:3.2.1.23) | 116.4 | 5.28 | 51 | 67.68 | - | Homotetramer | G | b.30.5.1; c.1.8.3; b.18.1.5; b.1.4.1 | Cytoplasmic | *lacZ (b0344)* | 0 | 10.4 |
| GL-$L_b$ | TIG_ECOLI | P0A850 | Trigger factor (EC:5.2.1.8) (TF) | 48.2 | 4.83 | 28 | 65.05 | One | Homodimer and monomer | O | i.1.1.2; d.241.2.1; d.26.1.1; a.223.1.1 | Cytoplasmic | *tig (b0436)* | 0 | 2.3 |
| | RPOA_ECOLI | P0A7Z4 | DNA-directed RNA polymerase subunit alpha (EC:2.7.7.6) (RNAP subunit alpha), RNA polymerase subunit alpha, Transcriptase subunit alpha | 36.5 | 4.97 | 15 | 56.53 | One | Homodimer | K | d.181.1.1; i.8.1.1; a.60.3.1; d.74.3.1 | Cytoplasmic | *rpoA (b3295)* | 1 | 4.0 |
| | PGK_ECOLI | P0A799 | Phosphoglycerate kinase (EC:2.7.2.3) | 41 | 5.08 | 22 | 73.9 | One | Monomer | G | c.86.1.1; c.1.1.1 | Cytoplasmic | *pgk (b2926)* | 1 | 2.5 |
| | OMPC_ECOLI | P06996 | Outer membrane protein C, Outer membrane protein 1B, Porin, OmpC | 40.3 | 4.48 | 20 | 77.38 | One | Homotrimer | M | f.4.3.1 | Outer membrane | *ompC (b2215)* | 0 | 9.7 |
| | OMPF_ECOLI | P02931 | Outer membrane protein F, Outer membrane protein 1A, Outer membrane protein B, Porin, OmpF | 39.3 | 4.64 | 24 | 82.6 | Two | Homotrimer | M | f.4.3.1 | Outer membrane | *ompF (b0929)* | 0 | 8.5 |
| | ALF_ECOLI | P0AB71 | Fructose-bisphosphate aldolase class II (EC 4.1.2.13) (FBP aldolase), Fructose-1,6-bisphosphate aldolase | 39.1 | 5.52 | 11 | 50.42 | Two | Homodimer | G | c.1.10.2 | Cytoplasmic | *fbaA (b2925)* | 1 | 7.2 |
| | GLF_ECOLI | P37747 | UDP-galactopyranose mutase (EC:5.4.99.9), UDP-GALP mutase, Uridine 5-diphosphate galactopyranose mutase | 43 | 6.61 | 27 | 79.02 | Five | Homodimer | M | d.16.1.7 c.4.1.3 | Cytoplasmic | *glf (b2036)* | 0 | - |
| | SUCC_ECOLI | P0A836 | Succinyl-CoA ligase [ADP-forming] subunit beta (EC:6.2.1.5), succinyl-CoA synthetase subunit beta | 41.3 | 5.37 | 22 | 76.8 | Five | Heterotetramer | C | c.23.4.1 d.142.1.4 | Cytoplasmic | *sucC (b0728)* | 0 | 6.7 |
| | MALE_ECOLI | P0AEX9 | Maltose-binding periplasmic protein, MBP, MMBP, Maltodextrin-binding protein | 43.3 | 5.22 | 13 | 51.77 | Five | - | G | c.94.1.1 | Periplasmic | *malE (b4034)* | 0 | - |
| | LACI_ECOLI | P03023 | Lactose operon repressor (LacI) | 38.5 | 6.39 | 24 | 80.28 | Five | Homotetramer | K | c.93.1.1; a.35.1.5 | Cyoplasmic | *lacI (b0345)* | 0 | 5.7 |
| | MANA_ECOLI | P00946 | Mannose-6-phosphate isomerase (EC:5.3.1.8), Phosphohexomutase, Phosphomannose isomerase (PMI) | 42.8 | 5.29 | 16 | 62.92 | - | Monomer | G | b.82.1.3 | Cytoplasmic | *manA (b1613)* | 0 | 3.6 |
| | TREC_ECOLI | P28904 | Trehalose-6-phosphate hydrolase (EC:3.2.1.93), Alphaalpha-phosphotrehalase | 63.8 | 5.51 | 31 | 62.61 | - | - | G | c.87.1.6 | Cytoplasmic | *treC (b4239)* | 0 | 4.3 |
| | AAT_ECOLI | P00509 | Aspartate aminotransferase (EC:2.6.1.1), AspAT, Transaminase A | 43.5 | 5.54 | 23 | 61.62 | Five | Homodimer | E | c.67.1.1 | Cytoplasmic | *aspC (b0928)* | 0 | 4.3 |

* GroEL substrate classes 1-3 are from Kerner et al. 2005, class 4 is from Fujiwara et al. 2010 and class 5 lists the proteins exclusive to Chapman et al 2006.

**Table S2** Flux Balance Analysis of GL-$L_b$ and GL-$H_b$ Strains.

| Strain | Metabolic Pathway | Metabolic Flux through the Pathway (mM/gm-DW/hr)* | | | |
|---|---|---|---|---|---|
| | | Flux in GL-$H_b$ | Flux in GL-$L_b$ | Flux Difference | Flux Ratio |
| GL-$H_b$ | Glycolysis/Gluconeogenesis | 289.2 | 59.6 | 229.6 | 2.3 |
| | Citric Acid Cycle | 183.9 | 100.1 | 83.7 | 0.9 |
| | Oxidative Phosphorylation | 103.3 | 48.9 | 54.5 | 1.1 |
| | Threonine and Lysine Metabolism | 41.2 | 0.6 | 40.6 | 6.0 |
| | Anaplerotic Reactions | 33.8 | 1.4 | 32.5 | 4.6 |
| | Inorganic Ion Transport and Metabolism | 62.7 | 31.6 | 31.1 | 1.0 |
| | Methylglyoxal Metabolism | 27.2 | 0.0 | 27.2 | NA |
| | Transport, Inner Membrane | 116.1 | 91.9 | 24.3 | 0.3 |
| | Glutamate Metabolism | 25.0 | 3.2 | 21.8 | 3.0 |
| | Alanine and Aspartate Metabolism | 126.3 | 111.6 | 14.8 | 0.2 |
| | Transport, Outer Membrane Porin | 28.1 | 17.0 | 11.1 | 0.7 |
| | Alternate Carbon Metabolism | 46.0 | 36.0 | 10.0 | 0.4 |
| | Glycine and Serine Metabolism | 9.8 | 0.8 | 9.0 | 3.6 |
| GL-$L_b$ | Membrane Lipid Metabolism | 0.4 | 0.4 | 0.0 | 0.0 |
| | Cofactor and Prosthetic Group Biosynthesis | 0.0 | 0.0 | 0.0 | -4.2 |
| | Nucleotide Salvage Pathway | 19.7 | 21.4 | -1.7 | -0.1 |
| | Pyruvate Metabolism | 452.6 | 478.4 | -25.8 | -0.1 |
| | Unassigned | 0.2 | 30.1 | -29.9 | -7.0 |
| | Pentose Phosphate Pathway | 218.9 | 266.8 | -47.8 | -0.3 |

* mM/gm-DW/hr – Millimolar Metabolite per Gram Dry Weight of the cell mass per hour.

**Table S2** Enriched Gene Ontology terms (level 3 - biological process terms), associated with the 404 proteins that were co-enriched/expressed with GroE across different experiments.

| Gene Ontology Terms | Protein Count | Fold Enrichment | *P*-value | Bonferroni Correction |
|---|---|---|---|---|
| GO:0006091: Generation of precursor metabolites and energy | 76 | 3.810 | 7.95e-24 | 1.28e-21 |
| GO:0044249: Cellular biosynthetic process | 193 | 1.541 | 5.72e-12 | 9.21e-10 |
| GO:0042180: Cellular ketone metabolic process | 84 | 2.130 | 4.30e-11 | 6.92e-09 |
| GO:0006082: Organic acid metabolic process | 82 | 2.118 | 1.04e-10 | 1.68e-08 |
| GO:0009308: Amine metabolic process | 73 | 2.003 | 1.55e-08 | 2.50e-06 |
| GO:0016052: Carbohydrate catabolic process | 42 | 2.525 | 9.87e-08 | 1.59e-05 |
| GO:0022900: Electron transport chain | 26 | 2.992 | 2.28e-06 | 3.68e-04 |
| GO:0006519: Cellular amino acid and derivative metabolic process | 58 | 1.908 | 3.06e-06 | 4.92e-04 |
| GO:0046483: Heterocycle metabolic process | 45 | 2.052 | 8.50e-06 | 1.37e-03 |
| GO:0006793: Phosphorus metabolic process | 29 | 2.405 | 3.27e-05 | 5.26e-03 |
| GO:0019538: Protein metabolic process | 60 | 1.672 | 9.90e-05 | 1.58e-02 |
| GO:0009059: Macromolecule biosynthetic process | 114 | 1.355 | 3.68e-04 | 5.76e-02 |
| GO:0006766: Vitamin metabolic process | 20 | 2.397 | 7.47e-04 | 1.13e-01 |
| GO:0006790: Sulphur metabolic process | 18 | 2.540 | 7.89e-04 | 1.19e-01 |
| GO:0016051: Carbohydrate biosynthetic process | 31 | 1.879 | 1.20e-03 | 1.75e-01 |
| GO:0044248: Cellular catabolic process | 29 | 1.864 | 2.06e-03 | 2.83e-01 |
| GO:0006461: Protein complex assembly | 11 | 3.214 | 2.28e-03 | 3.07e-01 |
| GO:0065003: Macromolecular complex assembly | 11 | 3.189 | 2.41e-03 | 3.22e-01 |
| GO:0005975: Carbohydrate metabolic process | 77 | 1.385 | 2.63e-03 | 3.46e-01 |
| GO:0033014: Tetrapyrrole biosynthetic process | 10 | 3.197 | 4.12e-03 | 4.86e-01 |
| GO:0044255: Cellular lipid metabolic process | 27 | 1.787 | 5.08e-03 | 5.60e-01 |
| GO:0009991: Response to extracellular stimulus | 10 | 3.016 | 6.02e-03 | 6.22e-01 |
| GO:0051186: Cofactor metabolic process | 30 | 1.640 | 9.63e-03 | 7.89e-01 |
| GO:0009057: Macromolecule catabolic process | 14 | 2.247 | 9.88e-03 | 7.98e-01 |

### Supplementary References.


Alexeeva, S., de Kort, B., Sawers, G., Hellingwerf, K.J., and de Mattos, M.J. (2000). Effects of limited aeration and of the ArcAB system on intermediary pyruvate catabolism in Escherichia coli. J Bacteriol *182*, 4934-4940.

Blais, E.M., Chavali, A.K., and Papin, J.A. (2013). Linking genome-scale metabolic modeling and genome annotation. Methods Mol Biol *985*, 61-83.

de Graef, M.R., Alexeeva, S., Snoep, J.L., and Teixeira de Mattos, M.J. (1999). The steady-state internal redox state (NADH/NAD) reflects the external redox state and is correlated with catabolic adaptation in Escherichia coli. J Bacteriol *181*, 2351-2357.

Faith, J.J., Driscoll, M.E., Fusaro, V.A., Cosgrove, E.J., Hayete, B., Juhn, F.S., Schneider, S.J., and Gardner, T.S. (2008). Many Microbe Microarrays Database: uniformly normalized Affymetrix compendia with structured experimental metadata. Nucleic Acids Res *36*, D866-870.

Fujiwara, K., and Taguchi, H. (2007). Filamentous morphology in GroE-depleted Escherichia coli induced by impaired folding of FtsE. J Bacteriol *189*, 5860-5866.

Kayser, A., Weber, J., Hecht, V., and Rinas, U. (2005). Metabolic flux analysis of Escherichia coli in glucose-limited continuous culture. I. Growth-rate-dependent metabolic efficiency at steady state. Microbiology *151*, 693-706.

Kirkpatrick, C., Maurer, L.M., Oyelakin, N.E., Yoncheva, Y.N., Maurer, R., and Slonczewski, J.L. (2001). Acetate and formate stress: opposite responses in the proteome of Escherichia coli. J Bacteriol *183*, 6466-6477.

McLennan, N., and Masters, M. (1998). GroE is vital for cell-wall synthesis. Nature *392*, 139.

Neidhardt, F.C., Ingraham, J.L., and Schaechter, M. (1990). Physiology of the Bacterial Cell: A Molecular Approach (Sinauer Associates).

Orth, J.D., Conrad, T.M., Na, J., Lerman, J.A., Nam, H., Feist, A.M., and Palsson, B.O. (2011). A comprehensive genome-scale reconstruction of Escherichia coli metabolism--2011. Mol Syst Biol *7*, 535.

Weber, J., Kayser, A., and Rinas, U. (2005). Metabolic flux analysis of Escherichia coli in glucose-limited continuous culture. II. Dynamic response to famine and feast, activation of the methylglyoxal pathway and oscillatory behaviour. Microbiology *151*, 707-716.

Wolfe, A.J. (2005). The acetate switch. Microbiol Mol Biol Rev *69*, 12-50.
